## Supplementary Material for "Liquid-liquid microphase separation leads to formation of membraneless organelles"

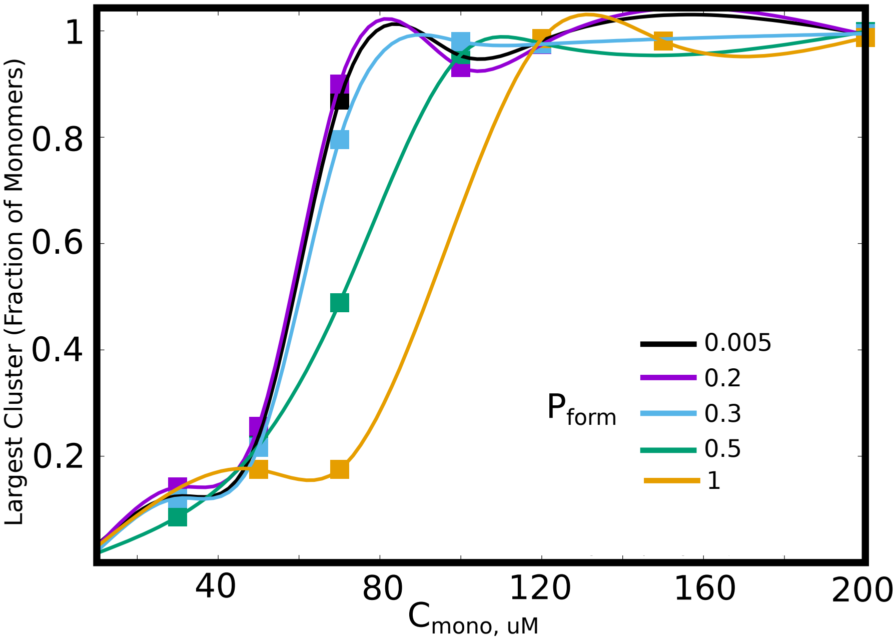


**Figure S1.** The size of the largest cluster, for different values of bond formation probability, $P_{form}$. A lower $P_{form}$ results in a slower arrest of the clusters, and thereby results in increased cluster sizes for smaller free monomer concentration, $C_{mono}$.

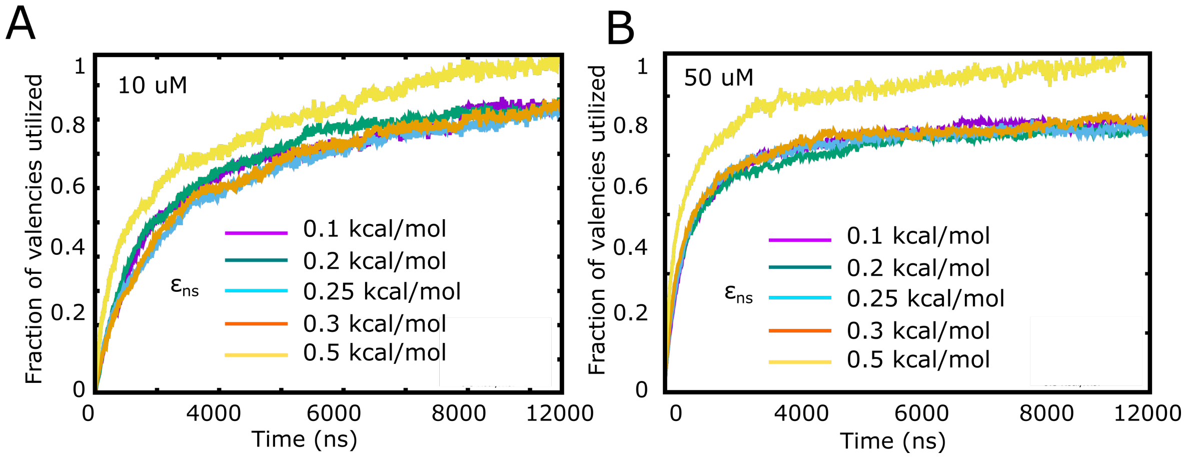


**Figure S2.** Fraction of valencies utilized as a function of increasing inter-linker interaction strength, for A) 10 $\mu$M and B) 50 $\mu$M free monomer concentration.

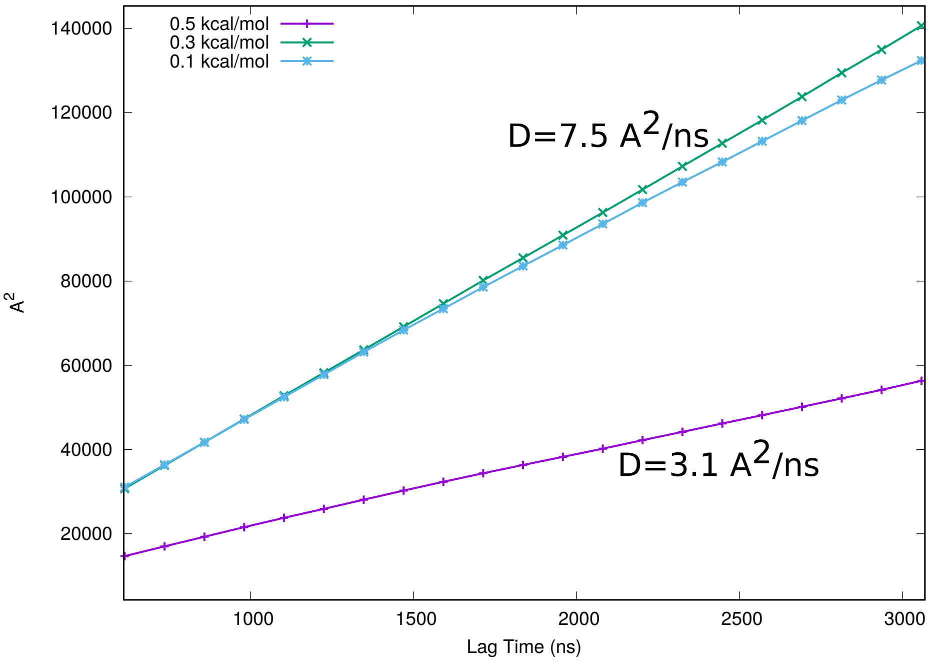


**Figure S3.** The mean-squared displacements of the center of masses of the constituent polymer chains as a function of increasing lag-times, in the absence of functional interaction. The three curves show the lagtimes for varying values of isotropic interaction strength between the linker residues.

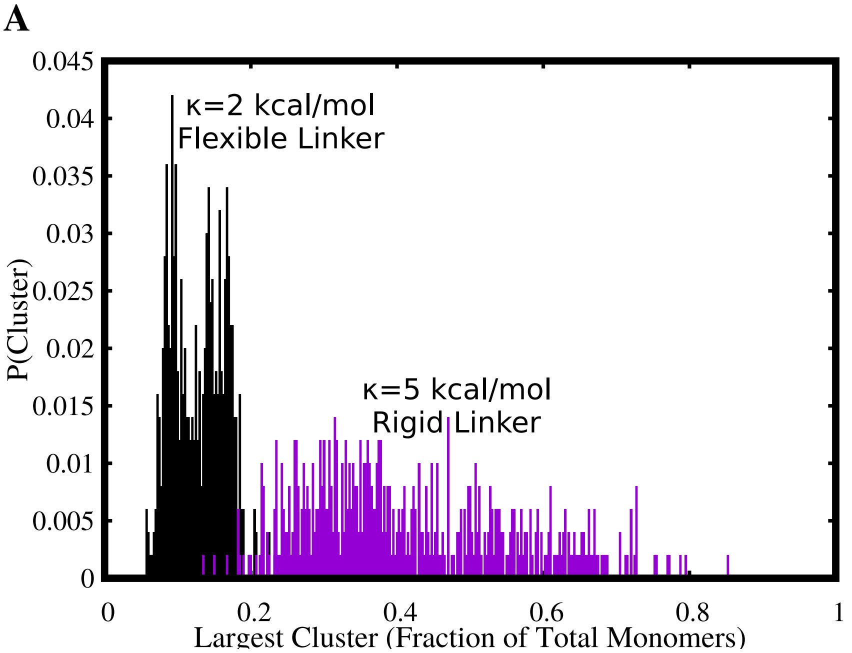


**Figure S4.** Comparision between the size distributions of the largest cluster, for stiff ($\kappa$=2 kcal/mol) versus flexible ($\kappa$=5 kcal/mol) linker regions. The free monomer concentration used for this plot was 50 $\mu$M and an a weak interlinker interaction strength of $\epsilon_n$ = 0.1 kcal/mol was used.

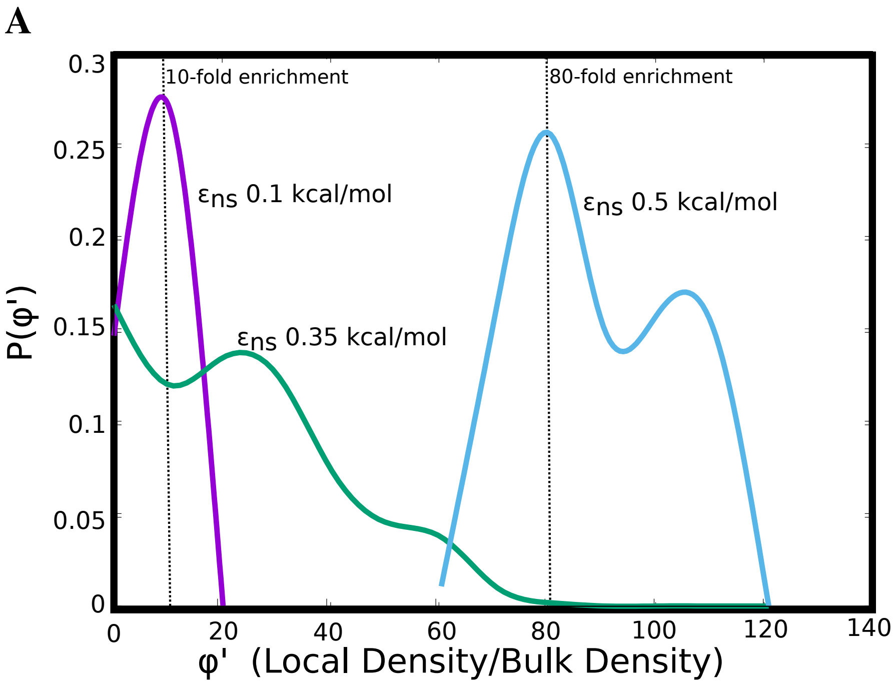


**Figure S5.** The probability of finding clusters with varying densities (normalized by the bulk densities) for different values of inter-linker interactions. As the inter-linker interactions increase, the degree of enrichment can go from 10-fold to 100-fold.

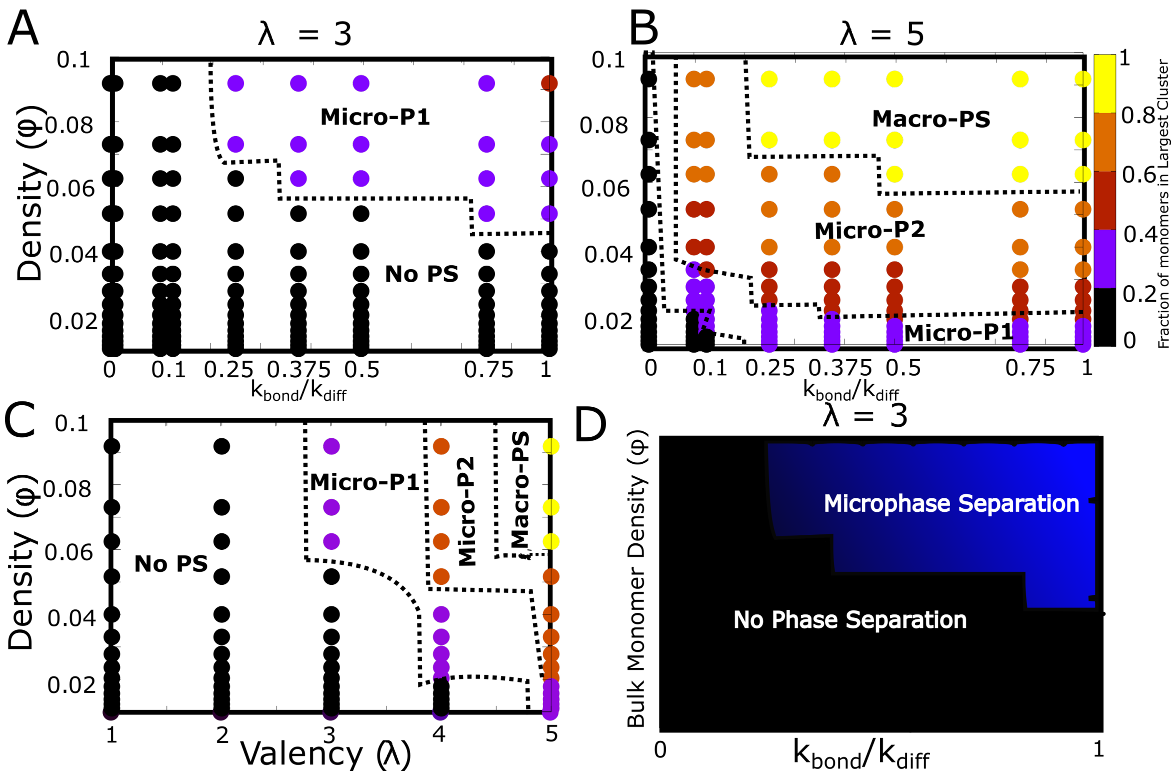


**Figure S6.** Detailed phase diagrams for A) and B) $\phi$-$k_bond$, C) $\phi$-$\lambda$ as the phase parameters. The cluster sizes were computed at the end of a simulation run of 2 hours (actual time), setting the rate of diffusion $k_diff$ to 1 $s^{-1}$. D) The bonding rate $k_{bond}$ was varied to identify the relationship between $k_{bond}$ and $k_diff$.

A

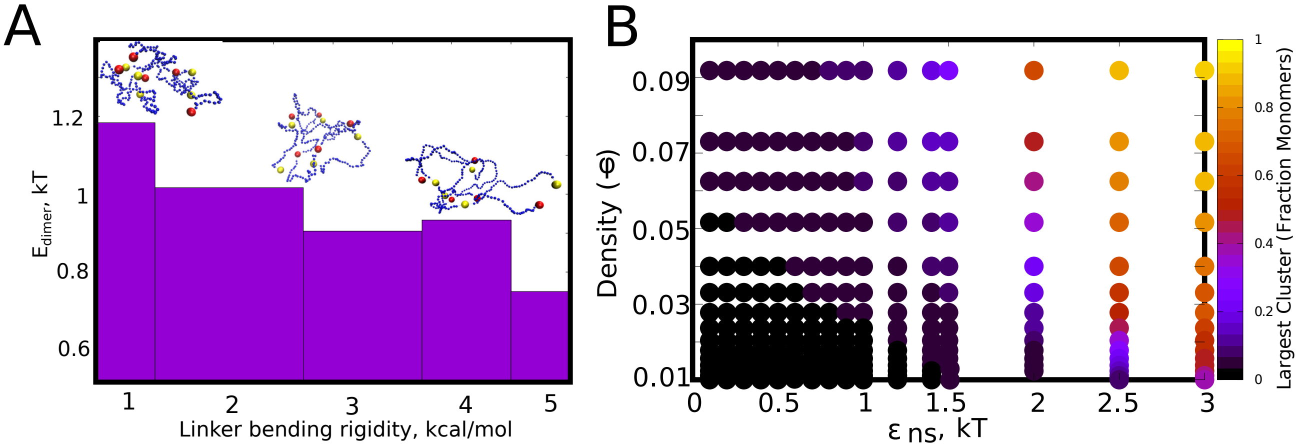


**Figure S7.** A) The mean pair-wise interaction energy for 100 different dimeric structures (from the LD simulations), for an inter-linker interaction strength of 0.1 kcal/mol, for different values of linker bending rigidity. B) The $\epsilon_{ns}$-$\phi$ phase diagram (for $\lambda$=0) showing no phase separation for low values of isotropic interaction strength. However, for values of $\epsilon_{ns} > 1$kT, phase separation is observed at the end of the simulation timescale of 2 hours (actual time).

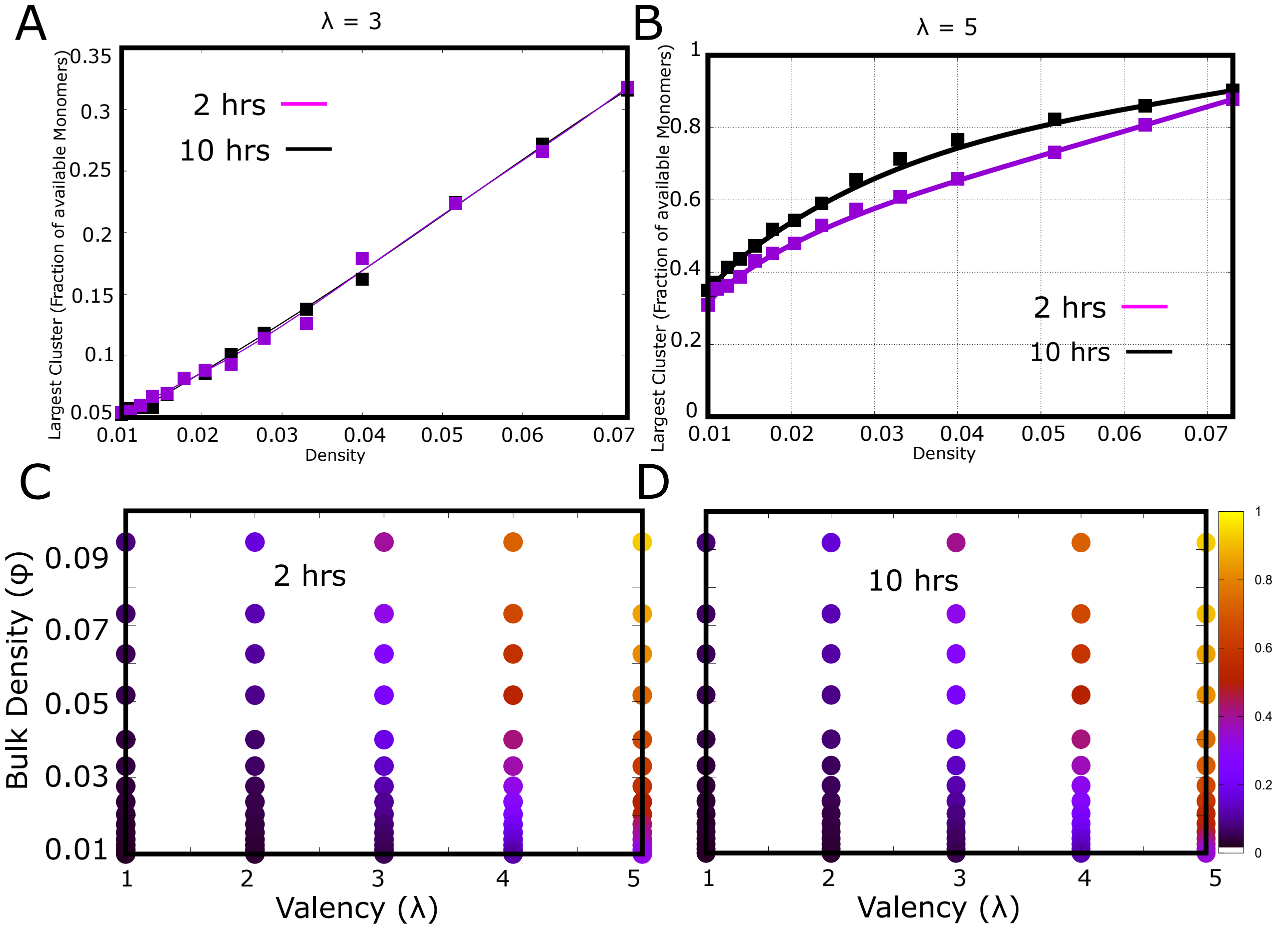


**Figure S8.** Convergence of phase diagrams. A) and B) shows the fraction of monomers in the largest cluster for 1 and 10 hours of actual time, for valency of 3 and 5, respecively. C) and D). The $\lambda$-$\phi$ phase diagram at the end of 2 and 10 hours of simulation time, respectively, showing very little difference.

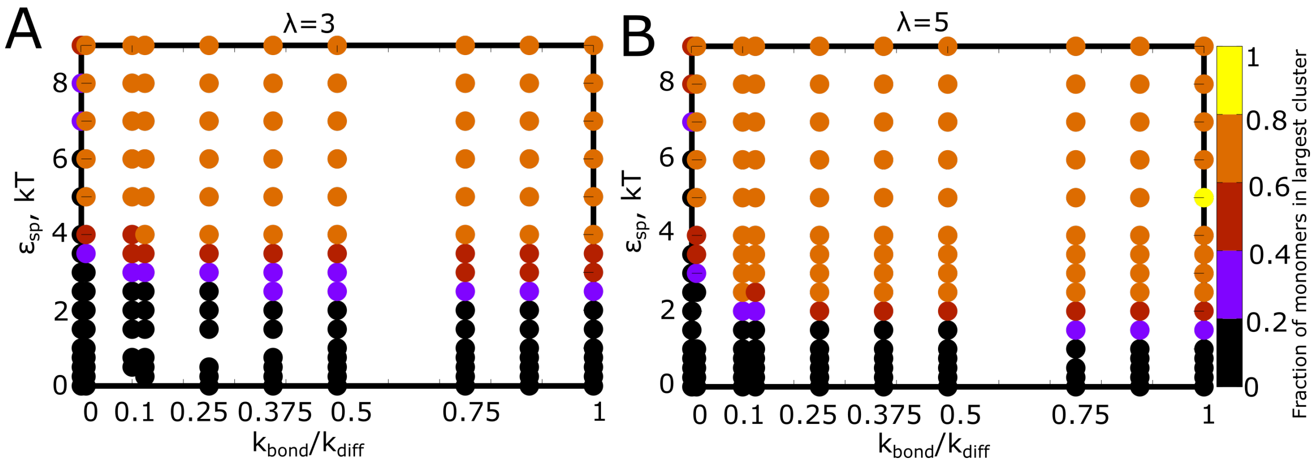


**Figure S9.** Detailed phase diagrams for A) and B) $\epsilon_{sp}$-$k_{bond}$ as the phase parameters. The bulk density of particles was set to 0.04, an intermediate density identified from the previous phase diagrams with density as a phase parameter. The cluster sizes were computed at the end of a simulation run of 2 hours (actual time), setting the rate of diffusion $k_diff$ to 1 $s^{-1}$.

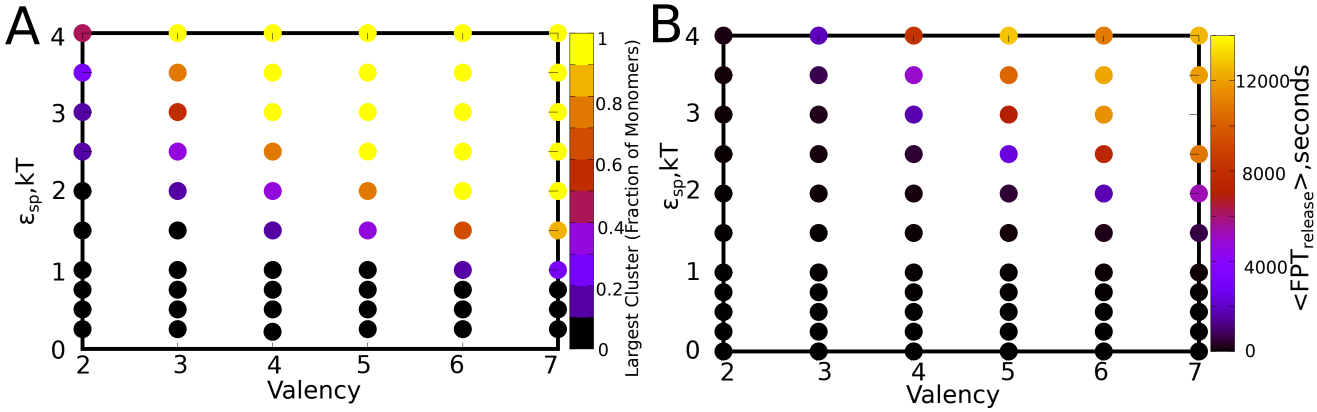


**Figure S10.** Mean cluster sizes for variation in $\epsilon_{sp}$ and $\lambda$, for a bulk density of 0.04, and a $k_{bond}/k_{diff}$ ratio of 1. B) Mean first passage times for a particle to exchange between a cluster and the bulk. The parameter values are same as in panel A.

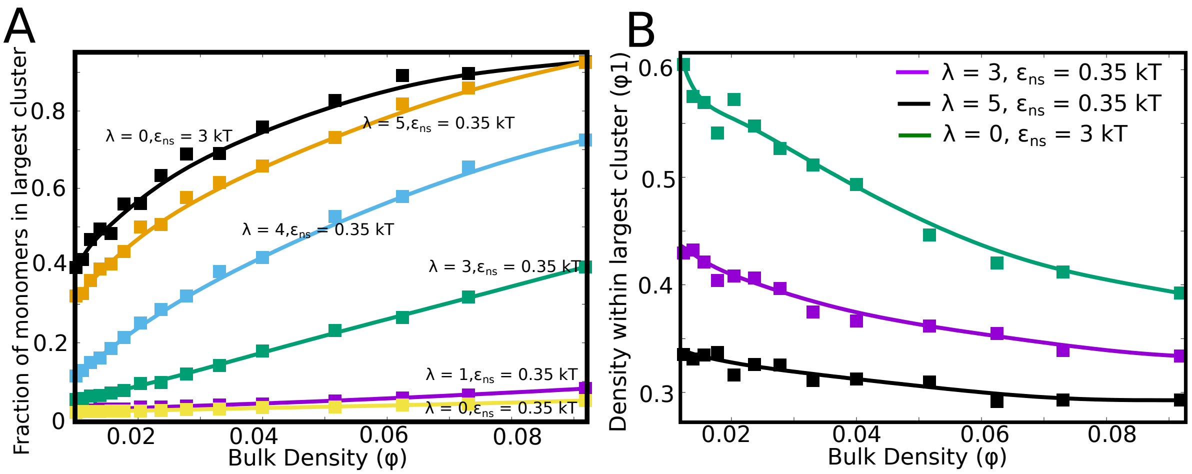


**Figure S11.** Cluster formation driven by isotropic versus specific interactions. A) Comparison of cluster sizes (as a fraction of total monomers) for different scenarios. The black curve shows cluster sizes for assembly driven by strong non-specific interactions alone ($\lambda$=0 and $\epsilon_{ns}$=3 kT). For a scenario involving weak isotropic interactions ($\epsilon_{ns}$ = 0.35 kT), the curves approach that of the isotropic interactions for higher valencies ($\lambda \to 4$). B) Densities of largest cluster for assemblies stabilized by isotropic interactions ($\epsilon_{ns}$ = 3kT, $\lambda$=0;green curve) and two different valencies (purple and black curves). Clusters stabilized by isotropic interactions are denser than the ones held together by specific interactions.
